# Metabolite toxicity as a driver of bacterial metabolite externalization

**DOI:** 10.64898/2026.01.17.700106

**Authors:** Ying-Chih Chuang, James B. McKinlay

**Affiliations:** Department of Biology, Indiana University, Bloomington, IN, USA; Biochemistry Program, Indiana University, Bloomington, IN, USA

## Abstract

Some microbes externalize costly biosynthetic precursors in sufficient quantities to sustain a recipient population through cross-feeding. However, it is unclear whether metabolites are externalized purely for a reciprocal benefit or if metabolite externalization also plays a physiological role for the producer. Here we focus on adenine, a metabolite externalized by some strains of the phototrophic bacterium *Rhodopseudomonas palustris* at sufficient levels to support *Escherichia coli* growth. In 10 long-term monocultures and 22 cocultures pairing *R. palustris* with *E. coli*, extracellular adenine externalized by all 140 isolates screened was 1.7 – 3.4-fold higher than that by the ancestor, suggesting that there was selective pressure for adenine externalization. We hypothesized that adenine is toxic to *R. palustris*. The CGA0092 growth rate decreased by half in the presence of about 0.3 mM external adenine. This inhibitory effect increased by an order of magnitude when we over-expressed adenine phosphoribosyltransferase to overcome a bottleneck in the purine salvage pathway, suggesting that toxicity stems from a metabolite derived from adenine. To assess whether adenine tolerance is connected to adenine externalization, we surveyed 12 evolved isolates and 49 environmental strains that externalized different levels of adenine, revealing a significant positive correlation. Our data suggests a physiological role for externalization of costly-metabolites like adenine at the origin of cross-feeding. In addition to cross-feeding, resulting metabolic interactions could be negative, considering that even a biosynthetic precursor like adenine can be inhibitory.

## INTRODUCTION

Microbial cross-feeding involves the transfer of a metabolite from a producer population to a recipient population. A prerequisite to cross-feeding is thought to be metabolite externalization (1), a term we use herein to encompass all possible externalization mechanisms (2). Some cases of metabolite externalization are easily explained, for example, where the metabolite is released by the producer as a waste product (3, 4). However, it is more difficult to explain why some producers externalize costly biosynthetic precursors, which at face value, might seem beneficial for the producer to retain.

Previously we discovered that the purple anoxygenic phototrophic bacterium *Rhodopseudomonas palustris* CGA0092 externalizes the purine nucleobase adenine (5, 6). Adenine externalization was accompanied by a high intracellular concentration relative to *R. palustris* TIE-1, which did not externalize adenine (6). We attributed this accumulation to a bottleneck in the purine salvage pathway; overexpressing the enzyme adenine phosphoribosyltransferase (Apt) eliminated externalization (6). We also surveyed 49 *R. palustris* strains and found that 16 strains externalized purines whereas the rest did not (6). This relatively high frequency of purine externalization across environmental isolates suggests that it is a relatively common trait for *R. palustris* strains, but the question remained as to why *R. palustris* would externalize a costly biosynthetic precursor.

Whereas it is possible that adenine externalization is reflective of a cross-feeding relationship in nature, adenine externalization might also play a physiological role. For example, cytoplasmic accumulation of intracellular metabolites can be sub-optimal for growth or even toxic (1, 2, 7-12). Maintaining homeostatic metabolite levels via regulating metabolic pathways is a commonly accepted mechanism. However, there is also growing acceptance that metabolite efflux plays a homeostatic role; efflux can ensure that metabolites are of sufficient availability to meet the demands of multiple pathways while avoiding harmful intracellular accumulation (1, 2, 7-12).

Here we test the hypothesis that purine externalization by *R. palustris* is associated with avoiding adenine toxicity. Support for this hypothesis stems in part from observations that long-term serial transfers of *R. palustris* CGA0092 derivatives universally enriched for strains that externalized more adenine than ancestral strains. Adenine was most toxic to *R. palustris* CGA0092 when purine salvage pathway activity was increased, suggesting that adenine toxicity is likely attributed to a downstream metabolite. Finally, we show a correlation between adenine externalization and resistance to environmental adenine in both evolved isolates and environmental strains. Our observations support a physiological role for metabolite efflux that could explain the origins of a spectrum of positive and negative metabolic interactions between bacteria.

## RESULTS

### Adenine externalization increased in long-term cultures

Some *R. palustris* strains like CGA0092 externalize adenine (6). We questioned whether adenine externalization is advantageous. To begin to address this question we examined three different *R. palustris* strains derived from CGA0092 that we had serially transferred in batch cultures for 134 - 503 generations in either monoculture or in coculture with either *E. coli* MG1655 or PFM2 (13, 14) (Fig 1A); none of the engineered mutations in these strains was responsible for adenine externalization (6). We predicted that if adenine privatization is advantageous then evolved isolates would externalize less adenine whereas if adenine externalization is advantageous, then adenine externalization would be maintained or elevated. We quantified extracellular adenine in supernatants from 3-6 evolved *R. palustris* isolates from each monoculture or coculture line (140 isolates in total). Each isolate was grown in monoculture, and then adenine was quantified in the supernatant using an *E. coli* purine auxotroph bioassay (6). All 140 evolved isolate supernatants had 1.7 – 3.4-fold higher adenine levels than in supernatants from the ancestral strains (Fig 1B). When comparing monoculture and coculture isolates that had both been transferred for ∼500 generations, average extracellular adenine levels were higher from evolved monoculture isolates versus coculture isolates (mean ± SD: 119 ± 12 vs 81 ± 16 μM/OD, respectively, p < 0.0001, unpaired t test with Welch’s correction, treating the average value from the 4 isolates from each line as a single replicate; n = 10). Mutations responsible for heightened adenine externalization were not obvious; mutations in nucleotide synthesis and salvaging genes were low and/or inconsistent in monocultures and cocultures (13, 14). We do not address responsible mutations herein, but the universally elevated adenine externalization implies that it was advantageous for *R. palustris* CGA0092 under the serial transfer conditions.

**Fig 1.**
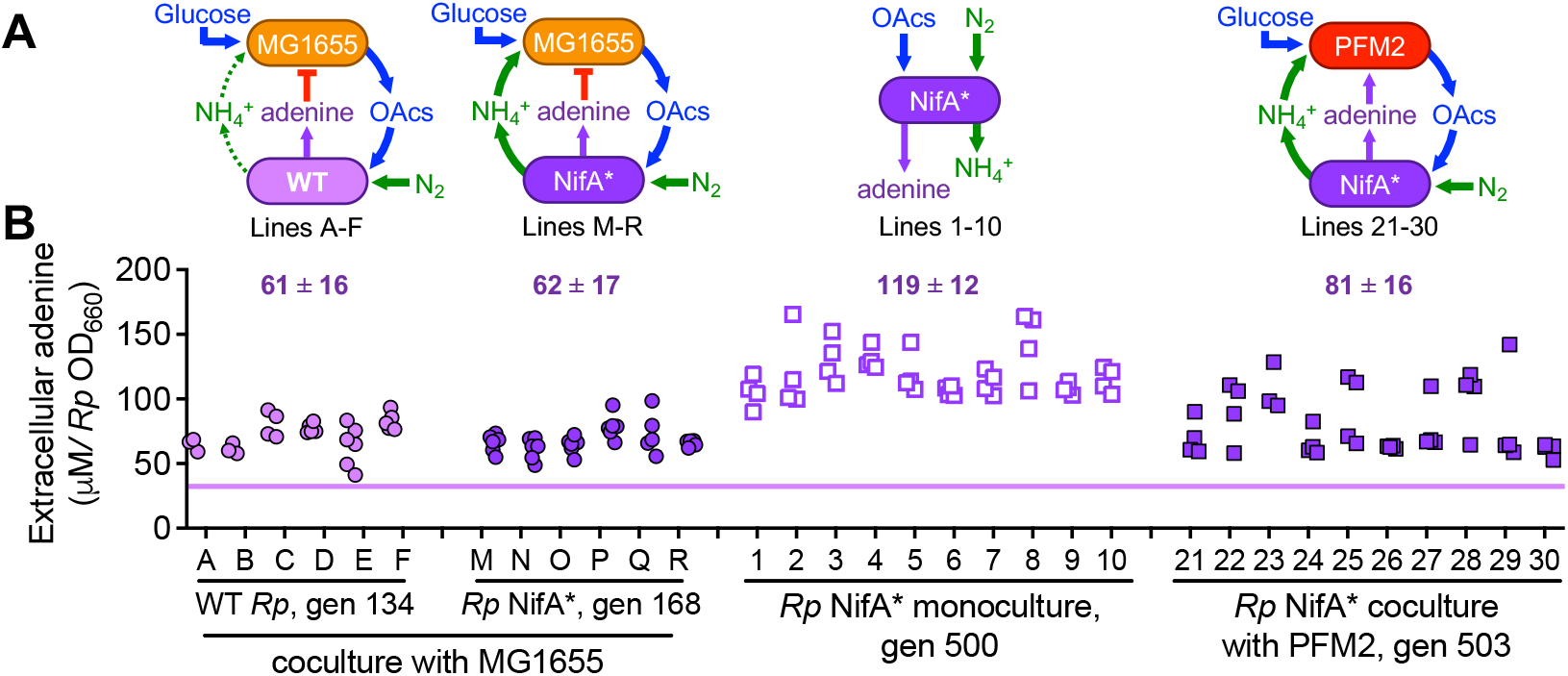
Adenine externalization increased in isolates evolved in various coculture or monoculture conditions. **A**. Coculture and monoculture conditions used for serial transfers in a minimal medium. Extracellular metabolites of relevance to this study are shown. *R. palustris* isolates from coculture lines A-F and M-R evolved from CGA4001 (Δ*hupS*) and CGA4003 (Δ*hupS, nifA*\*), respectively. *R. palustris* isolates from monoculture lines 1-10 and coculture lines 21-30 evolved from CGA676 (*nifA*\*). **B**. Adenine measured using the *E. coli ΔpurH* bioassay with supernatants from evolved isolates grown in monoculture in a minimal medium (n=4-6). Each point represents an individual isolate. The horizontal purple line is the range (thickness) of extracellular adenine for CGA676 and CGA4003 combined. Numerical values show the mean extracellular adenine ± SD for a given condition by treating the mean of for all isolates for a given evolved line as single value; n= 6 or 10.

### Extracellular adenine can be toxic to *R. palustris* CGA0092

Metabolite externalization can be a homeostatic mechanism to prevent cytoplasmic metabolites from reaching toxic levels (1, 2, 7-12). We thus hypothesized that adenine externalization could be associated with avoiding toxic accumulation of adenine or related metabolites. Adenine externalization by CGA0092 was previously attributed to a bottleneck in the purine salvage pathway caused by low expression of *apt*, encoding phoshporibosyltranserase that converts adenine to AMP; over-expressing *apt* eliminated adenine externalization (6) (Fig 2A). We predicted that if low *apt* expression helps protect the cell by contributing to adenine externalization, then overexpressing *apt* should be detrimental.

**Fig 2.**
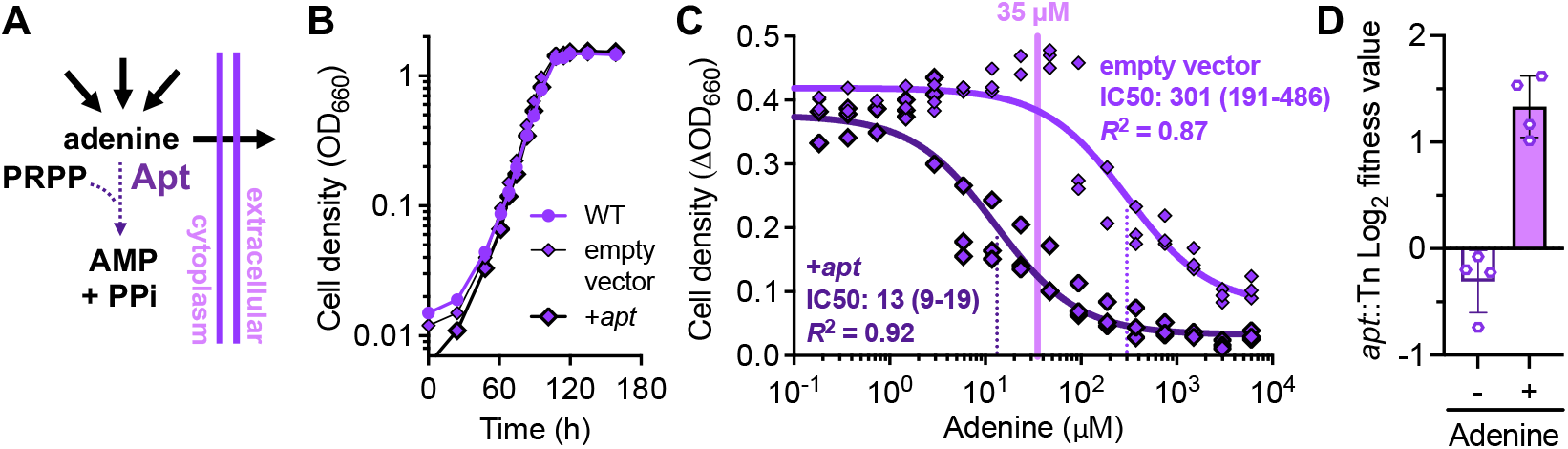
Higher Apt expression leads to increased adenine sensitivity. **A**. Various pathways generate adenine that can then be converted to AMP via adenine phosphoribosyltransferase (Apt) or externalized. Adenine externalization was previously found to correlate with low expression of Apt. **B**. Representative *R. palustris* monoculture growth curves in a minimal medium in test tubes. Similar trends were seen for three biological replicates without (empty vector, pBBPgdh) and with (+*apt*, pBBPgdh-apt) expression of an additional copy of *apt* from a plasmid. WT, CGA0092. **C**. Effect of supplemented adenine on *R. palustris* CGA0092 growth in a minimal medium in 96-well plates without and with *apt* expression from a plasmid. IC50 values (dotted lines; μM with non-symmetrical 95% CI) were determined by non-linear regression using the Prism 10 ‘[inhibitor] vs response (3 parameters)’ function. The solid purple bar is the adenine concentration in CGA0092 supernatants at an OD_660_ of 1 (∼35 µM). Each point represents an individual culture. **D**. Log_2_ fitness values for CGA009 mutants with a transposon disruption of *apt* as part of a TnSeq experiment. Mutant libraries were grown phototrophically in minimal photosynthetic medium with 10 mM butyrate, 10 mM NaHCO_3_, without or with 2 mM adenine. Fitness values are based on the change in mutant frequency from the time of inoculation, relative to all other mutants in the pool. Each point represents a replicate TnSeq experiment.

To test this prediction, we expressed *apt* from a plasmid in CGA0092. Over-expression of *apt* had no effect on culture growth compared to an empty vector control (Fig. 2B). However, we considered that high *apt* expression might only be detrimental in the presence of exogenous adenine. This scenario would be analogous to the emergence of an *apt*-expressing mutant, surrounded by a large population of adenine-externalizing relatives. Indeed, when the media was supplemented with different levels of adenine the IC50 value for the empty vector control strain was ∼301 μM whereas the IC50 value was 13 μM for the *apt*-expressing strain (Fig 2C). In support of these observations, we also observed that transposon disruption of *apt* in CGA009 (exhibits similar adenine externalization levels as CGA0092 (6)) led to a 2 – 3-fold increase in fitness in the presence of 2 mM adenine compared to a pool of genome-wide transposon mutants in a TnSeq experiment (Oda et al. in prep) (Fig 2D). In comparison, disruption of *apt* in the absence of adenine had a negligible or slightly negative effect on fitness (Fig 2D).

### Purine externalization is correlated with adenine tolerance

With evidence that extracellular adenine can be toxic to CGA0092, we hypothesized that the elevated adenine externalization observed in evolved isolates (Fig 1) improved adenine tolerance. To test this hypothesis, we compared how an adenine supplement affected the growth rates of ancestral strains versus evolved isolates. Since we previously observed that all 140 evolved isolates showed elevated adenine externalization (Fig 1), we felt it was redundant to examine all lines and settled on examining a single isolate from three evolved lines from each serial transfer condition (12 isolates in total).

All of the ancestor replicates were negatively impacted by 0.3 mM adenine; growth rates were 44 – 86% of that of replicates without an adenine supplement (Fig 3). Nine of the twelve evolved isolates tested were also negatively impacted by adenine. However, the impact was less severe, with growth rates ranging from 73 – 90% of those without an adenine supplement. The other three isolates showed higher growth rates with adenine, ranging from 101 – 123% of those without an adenine supplement. Overall, there was correlation between adenine externalization level and resistance to adenine; linear regression was poor (R^2^, 0.13) but the nonparametric Spearman correlation was significant (p = 0.01).

**Fig 3.**
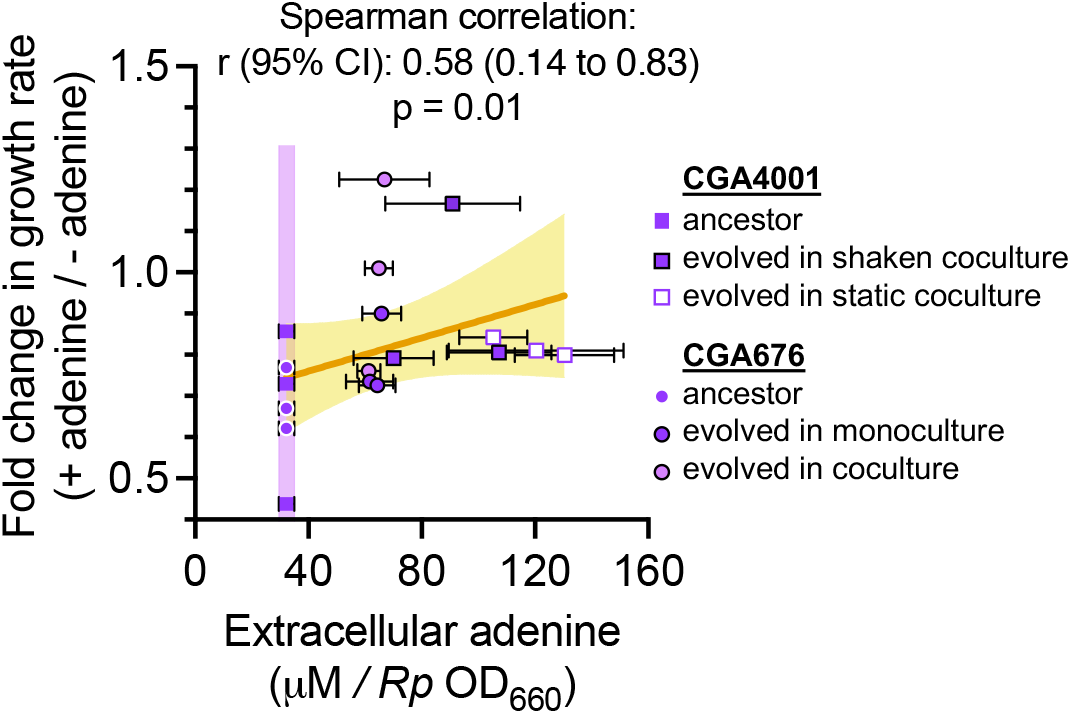
Correlation between adenine externalization and adenine resistance in evolved isolates. Growth rates were compared in MDC with 10 mM acetate, 10 mM NH_4_Cl, anoxic test tubes with and without 0.3 mM adenine. Each data point represents the growth rate for a given culture: three replicates of each ancestor strain or a single evolved isolate from lineages A, B, E, M, N, O (derived from the CGA4001 ancestor, squares) and lineages 1, 2, 3, 21, 22, 23 (derived from the CGA676 ancestor, circles). The purple vertical bar highlights the ancestral range of adenine externalization. The orange line is the linear regression of all data points (y = 0.002x + 0.679; R^2^, 0.13) ± 95% CI in yellow. Spearman correlation analysis assumed a nonparametric distribution.

Intrigued by a possible correlation between adenine externalization and tolerance, we hypothesized that this trend might be more pronounced for environmental *R. palustris* isolates that exhibit a wider range of purine externalization levels: undetectable to > 150 µM per OD_660_ by our bioassay that does not distinguish purines (6). To test this hypothesis, we measured culture growth for 49 *R. palustris* strains grown in 96-well plates under phototrophic conditions with different levels of added adenine. Due to the high-throughput format and the requirement for both light and anaerobic conditions, we could not monitor growth rates in a plate reader but instead measured culture turbidity at a 6-day time point where inhibited cultures might still be growing slowly or not at all. As predicted, there was good correlation between purine externalization and adenine tolerance based on whether a culture achieved 50% of the final cell density (OD_660_) observed in the 0.0002 mM adenine condition; the Spearman correlation was 0.67 with a significant p value of <0.0001. Strains that externalized 40 - 170 µM purine could tolerate the addition of 0.2 – 2 mM adenine, whereas strains that externalized less purine tolerated 0.02 to 0.2 mM adenine, with a few exceptions (Fig 4). CGA009 tolerated more adenine than expected in this experiment when compared to the IC50 assay (Fig 4 vs Fig 2). The discrepancy could be due to the age of the adenine stock or variable adenine degradation between stock solutions resulting from necessary sonication to dissolve adenine (15). Regardless, the general correlation between adenine externalization and tolerance was clear. Our data suggests that adenine externalization is a strategy used by diverse, but not all, *R. palustris* strains to tolerate toxic aspects of adenine.

**Fig 4.**
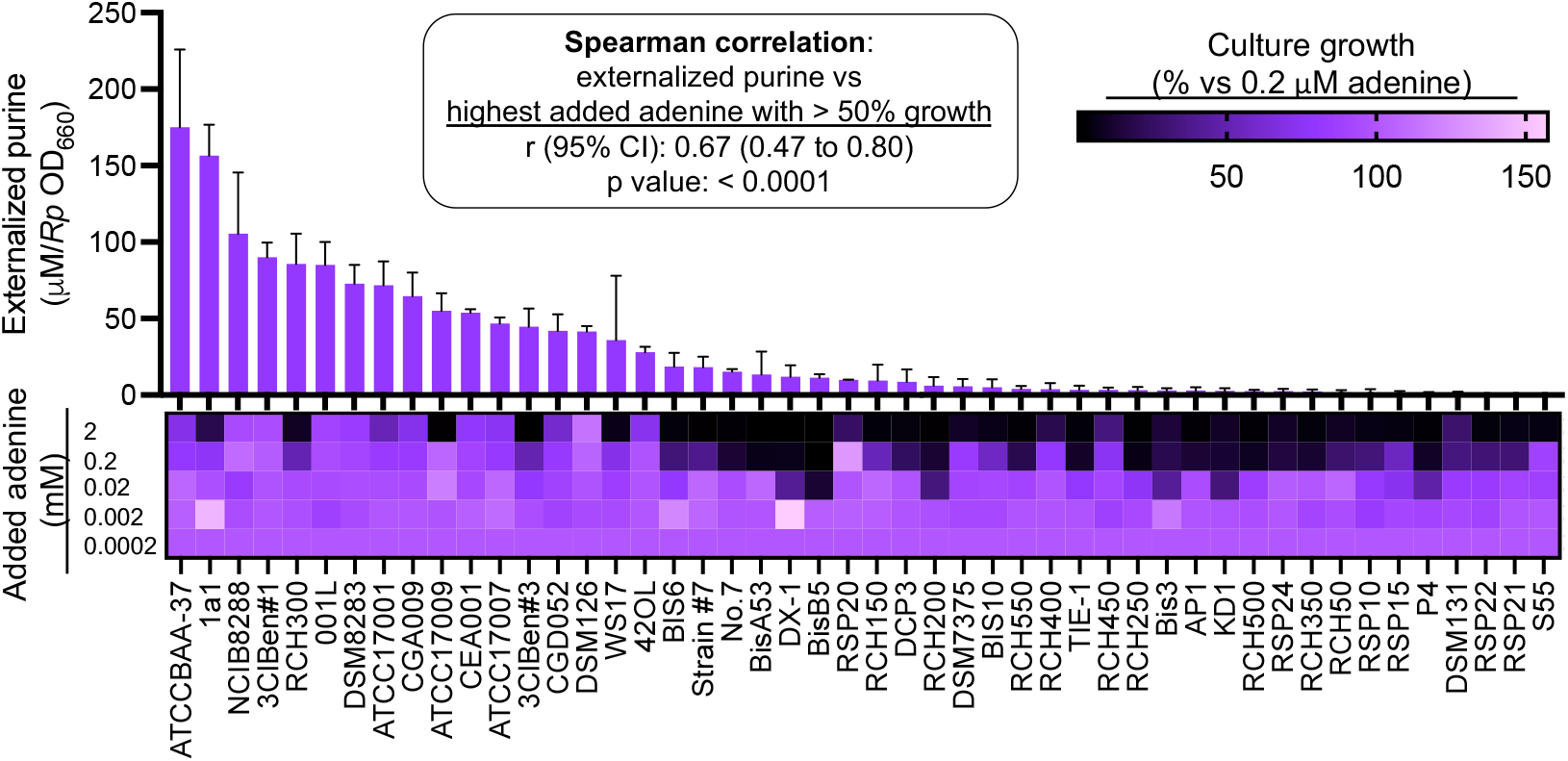
Purine-excreting *R. palustris* strains tend to tolerate more adenine. Externalized purine data (top graph) is reproduced from published values (6); error bars = SD; n = 3. Strain phylogeny and source locations are published elsewhere (6). Heatmap (bottom graph) shows percent final growth relative to growth with 0.2 µM adenine. All cultures were grown in MDC with 10 mM sodium acetate and 10 mM NH_4_Cl in a 96-well plate using a 1% inoculum from starter cultures grown in similar conditions without adenine. Culture growth was determined by turbidity (OD_660_) after 6 days. A single culture was grown for each of the 5 adenine concentrations. Spearman correlation analysis assumed a nonparametric distribution and compared the amount of adenine externalized to the highest level of adenine supplement where > 50% of the final cell density (OD_660_) of the 0.0002 mM condition was observed.

## DISCUSSION

We found that adenine externalization and tolerance are correlated in *R. palustris*. This correlation was observed for isolates that had evolved greater levels of adenine externalization through serial transfers (Fig 3) and a stronger correlation was observed for environmental isolates (Fig 4). We reason that adenine externalization avoids accumulation of intracellular adenine to toxic levels. Intracellular adenine levels are high in adenine-externalizing CGA0092 compared to TIE-1, which does not externalize adenine (6). Adenine itself might not be toxic, but rather a derivative of adenine; forcing excess adenine into the purine salvage pathway by over-expressing *apt* increased adenine sensitivity whereas *apt* transposon mutants had elevated fitness in the presence of adenine (Fig 2). Adenine externalization by *R. palustris* might be analogous to a mechanism used by *E. coli* to control pyrimidine levels, where accumulation of uridine and cytidine phosphates were avoided by degradation to uracil, followed by uracil externalization (16).

The increased adenine externalization observed by all 140 evolved isolates analyzed (Fig 1) suggests that adenine externalization was still under selection, at least under our lab conditions. It is possible that batch culture conditions allowed adenine to accumulate to levels that selected for strains with higher adenine externalization. In natural environments, one might anticipate greater dilution or diffusion of adenine from the local environment, or removal by neighboring cells. In support of this view, *R. palustris* isolates that evolved in coculture with *E. coli* exhibtied lower adenine externalization than isolates that evolved in monoculture (Fig 1), possibly because consumption of adenine by *E. coli* helped alleviate toxicity. There is good evidence that *E. coli* uses externalized adenine: (i) it down-regulates purine synthesis in response to added adenine and (ii) when in coculture with *R. palustris* where adenine is produced (14), (iii) *R. palustris* CGA0092 derivatives rescue *E. coli* purine auxotrophs in coculture, and (iv) *R. palustris* CGA0092 supernatants support *E. coli* purine auxotroph growth (17).

The mode of adenine externalization, and the mutations that enabled greater adenine externalization, remain unclear. Previously, a computational modeling approach suggested that passive diffusion across the membrane could explain extracellular levels (6). However, intracellular and extracellular concentrations did not equilibrate in stationary phase, suggesting the possibility of controlled transport, for example by efflux proteins (6). Evolution of higher adenine externalization might also suggest involvement of an efflux protein or perhaps less re-uptake by another transporter; metabolite externalization is often the sum of opposing efflux and uptake (10, 12). Currently, we also cannot rule out changes to membrane composition that might influence the extracellular adenine level. Unfortunately, mutations in evolved isolates do not give a clear indication of responsible mutations (14).

Our results suggest that adenine externalization has a physiological role, but we know that this can also set the stage for metabolic interactions. We previously found that the level of externalized adenine is sufficient to support the growth of an *E. coli* purine auxotroph population in coculture (6). In certain conditions, obligate reciprocal cross-feeding can easily be established; we used conditions where *R. palustris* was reciprocally dependent on *E. coli* for carbon in the form of fermented organic acids (6). Thus, as has been postulated by others, the prerequisite of metabolite externalization for cross-feeding might often stem from physiological roles, such as the externalization of biosynthetic precursors to maintain homeostatic intracellular levels (1, 2, 8, 9, 12)

In different contexts, externalization of the same metabolites can instead be inhibitory. For example, fermentation products, which can be used as a carbon source by neighbors, can also inhibit growth by lowering the pH (18, 19). Given our observations herein that extracellular adenine can be inhibitory to some *R. palustris* strains, one must also consider negative interactions stemming from externalized biosynthetic precursors. Separately we observed that some mutations enriched in *E. coli* MG1655 in coculture with *R. palustris* were consistent with growth-inhibition by adenine (14) (Fig 1A). The potential for an externalized nucleobase to benefit or harm neighboring cells goes beyond domesticated *E. coli*. A recent study that screened nearly 7,000 bacterial isolates from 27 environmental soil samples found that most samples contained nucleotide auxotrophs (20), and thus would benefit from externalized nucleobases. However, the same study later excluded nucleobases from the growth medium due to an inhibitory effect on many of the isolates (20). Thus, we posit that homeostatic externalization of biosynthetic precursors sets the stage for broad metabolic interactions that can include a range of positive or negative effects on a recipient.

## MATERIALS AND METHODS

### Bacterial strains and plasmids

*R. palustris* strains CGA676, CGA4001, and CGA4003 were derived from CGA0092 (21). Strains with *nifA*\* mutations (CGA676 and CGA4003) excrete NH_4_^+^ under N_2_-fixing conditions (18, 22). Strains with Δ*hupS* (CGA4001 and CGA4003) are incapable of H_2_ oxidation. Evolved isolates were always compared to their respective *nifA*\* ancestor. In all other cases, the type strain CGA0092 was used. Information on all other strains, including CGA0092 harboring pBBPgdh (empty vector) (22) and pBBPgdh-apt (+*apt*), and environmental isolates can be found elsewhere (6, 13).

### Growth conditions

To culture *R. palustris* in test tubes, anoxic media were prepared by bubbling N_2_ through 10 ml of minimal M9-derived coculture medium (MDC) (18) in 27-ml anaerobic test tubes, then sealing with rubber stoppers and aluminum crimps prior to autoclaving. After autoclaving, MDC was supplemented with 20 mM sodium acetate and 10 mM NH_4_Cl (final concentrations) via syringe from anoxic stock solutions. Plasmid-carrying strains were also supplemented with 100 μg ml^-1^ kanamycin (km). All starter cultures were inoculated with single colonies from agar plates. Test cultures were inoculated with a 1% v/v inoculum of starter culture. For the TnSeq experiment, cultures were grown in anaerobic test tubes in minimal photosynthetic medium (PM) (23, 24), with 10 mM sodium butyrate and 10 mM sodium bicarbonate without or with 2 mM adenine as described (Oda et al. in prep). All cultures were grown at 30°C with light from a 45 W halogen bulb (430 lumens) and shaking at 150 rpm; tubes were horizontal.

For cultures grown in 96-well plates, each well contained 0.2 ml of MDC, prepared using oxic stock solutions and inoculated under oxic conditions. Plates were then sealed inside a clear plastic anerobic latch-box (BD) and anoxic conditions were established using BD GasPak EZ sachets.

### Analytical procedures

Cell densities were measured via turbidity (OD_660_) using a Genesys 20 visible spectrophotometer (test tubes) or a BioTek Synergy H1 plate reader (96-well plates). To measure extracellular purines (assumed to be adenine), evolved cocultures and monoculture frozen stocks (25% glycerol) were struck for colony isolation on PM agar with 10 mM succinate and incubated anaerobically at 30°C with light. Randomly selected *R. palustris* isolates were then grown as monocultures in MDC with acetate and NH_4_Cl as above. Adenine was then quantified in stationary phase supernatants using an *E. coli* Δ*purH* bioassay, based on a linear relationship between final culture turbidity and adenine concentration (6). Methodology for the transposon sequencing experiment and determination of fitness values is described elsewhere (Oda et al. in prep).

### Statistical analyses

Statistical analyses were performed using Graphpad Prism (v10).

## ACKNOWLEDGEMENTS

This work was supported in part by US Army Research Office grants W911NF-14-1-0411 and W911NF-17-1-0159, the National Science Foundation CAREER award MCB-1749489.

We thank A. Dalia, J. Drummond, J.P. Gerdt, M. Federle, and the McKinlay lab for helpful comments.

